# Evolutionarily conserved clusters of colon with lung cancer susceptibility loci, linked with most *DUSP* phosphatase genes, may help to dissect mechanisms of cancer susceptibility

**DOI:** 10.1101/2021.10.13.461047

**Authors:** Lei Quan, Miranda Lynch, Peter Demant

## Abstract

We show evolutionarily conserved pairwise genetic linkage and clustering of majority of colon and lung cancer susceptibility QTLs in mice, rats and humans. The patterns of susceptibility or resistance to these two cancers in recombinant congenic mouse strains were concordant and the responsible susceptibility loci closely linked, in spite of completely different carcinogens and protocols used for induction of the two tumors. Most *DUSP* (Dual specificity phosphatase) genes are linked to these clusters. These data suggest that an important part of colon and lung cancer susceptibility is controlled by related and evolutionarily conserved processes.

---

Family studies and animal experiments have shown that whereas the familial cancers comprising less than 10% of cases are caused by germ-line mutations of high penetrance cancer genes, the common cancers are under control of multiple susceptibility genes^1,2,3^ with limited effects. Subsequent genome wide associacion studies (GWAS) in humans have indicated map positions of many of these genes^4^, but their identity and functions have not yet been sufficiently clarified. We show here in mice, rats, and humans that most colon and lung cancer susceptibility loci exhibit four unexpected genetic characteristics: a. They are mostly linked pairwise together in genome b. Their linkage and clustering in multiple short chromosomal regions are evolutionarily conserved for >70 million years c. Their effects on susceptibility to colon and lung tumors in recombinant congenic strains are highly correlated d. Most *DUSP* (*Dual specificity phosphatase*) genes are significantly and in many ilinstances tightly linked with these colon-lung cancer susceptibility gene clusters as well as with susceptibility genes for other cancers. A comparative study of effects of these clustered genes in different species and organs may clarify the mechanisms of their action in tumorigenesis.

## 1. Co-mapping of colon and lung cancer susceptibility loci

Before the GWAS technologies became broadly used, we have mapped in mice numerous susceptibility genes for colon and lung cancer using recombinant congenic (RC) strains^3^. These linkage studies^5^ (Fig. 1A) showed that 12 of the 15 detected colon cancer susceptibility loci (mostly used locus symbol *Scc* - *Susceptibility to colon cancer*) and 22 of the published 41 lung cancer susceptibility loci (mostly used locus symbols *Sluc* – *Susceptibility to lung cancer* or *Pas* – *Pulmonary adenoma susceptibility)* are frequently pair-wise linked together^5^, forming 19 clusters (14 of them shorter than 2.5 cM) that included also 10 pre-GWAS human and 3 rat^6^ colon cancer susceptibility loci, each cluster containing one or more colon as well as one or more lung cancer susceptibility locus (Fig. 1B).

**Figure 1.**
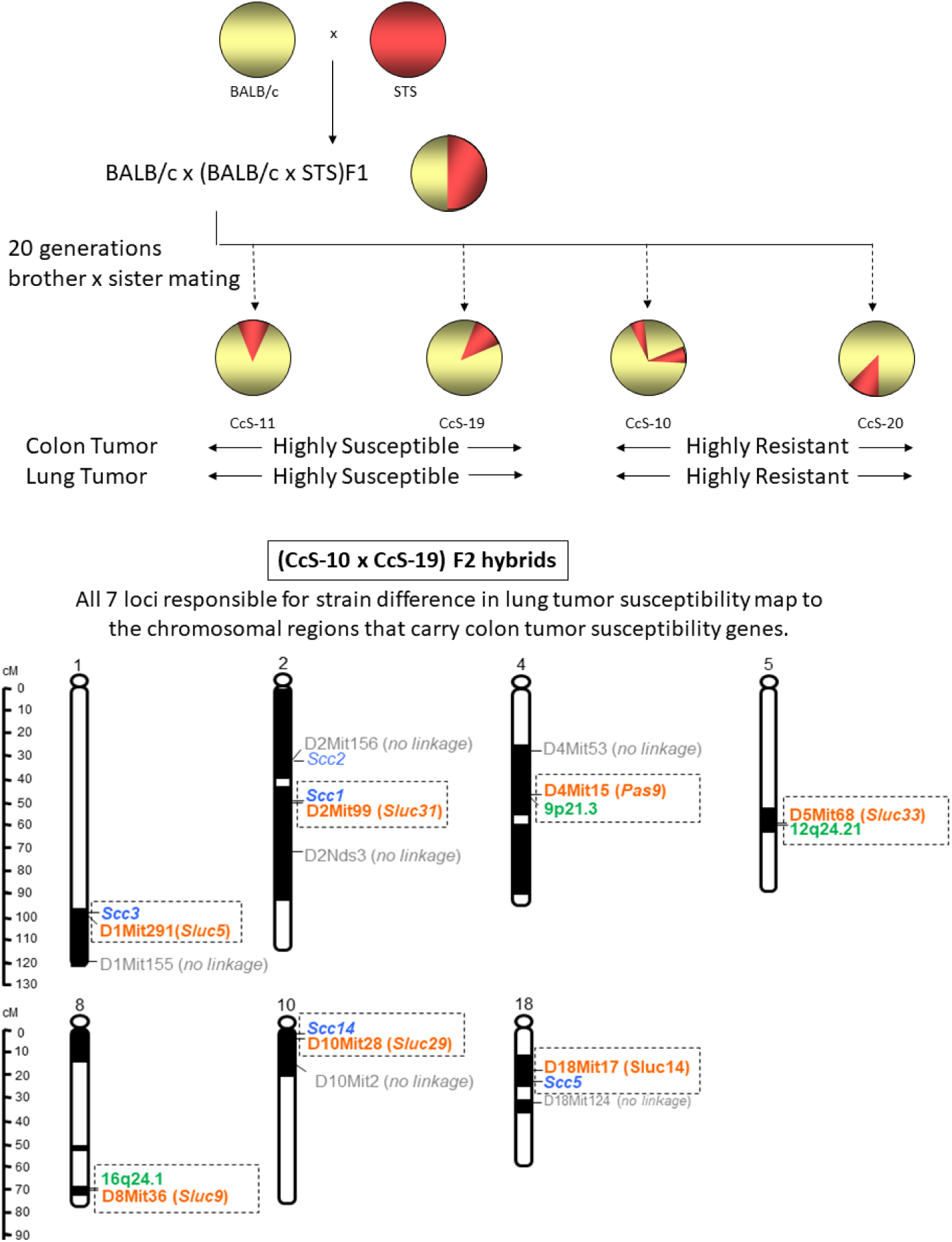
Concordant susceptibility of recombinant congenic (RC) strains to colon and lung tumors and co-localization of the responsible loci. A schematic representation of the derivation of recombinant congenic strains by random backcrossing of STS and BALB/c mice and subsequent inbreeding. The resulting RC strains carried each a different set of ± 12.5 % STS-derived genes on BALB/c genetic background. These sets overlapped only minimaly with the STS-derived sets in other RC strains. Two RC strains, CcS-10 and CcS-20, were highly resistant to both colon and lung tumors, whereas strains CcS-11 and CcS-19 were highly susceptible to both tumors. Mapping of <u>lung</u> tumor susceptibility loci controlling difference between CcS-10 and CcS-19 in (CcS-10 x CcS-19)F2 hybrids revealed 7 loci, all of which were closely linked to chromosomal locations of mouse or human <u>colon</u> tumor susceptibility loci.

This genetic clustering was obvious in spite of different species, induction protocols, and in spite of mouse colon and lung tumors being analyzed in crosses of different strains, mostly BALB/c and STS/A for colon tumors, and mainly O20/A and C57BL/10 or A/J and C57BL/6 for lung tumors^3,5^, and by different carcinogens: colon cancers by 1,2 dimethyl-hydrazine (DMH), or azoxymethane (AOM) ^5,7,8^ and lung cancers by N-ethyl-N-nitroso-urea (ENU), or ethylcarbamate (urethane) ^5,9^ (Fig. 1C).

## 2. GWAS data confirmed the intra- and inter-species co-localization of colon and lung cancer susceptibility loci

New GWAS data based on the reliable standard p-value threshold of 5 × 10^−8^ revealed 31 novel colon cancer susceptibility loci in humans^10^. These data conformed with our previous report^5^ of evolutionary conservation of co-localization of colon and lung cancer susceptibility^11^. We projected the map positions of these 31 human colon cancer susceptibility loci on their orthologous locations on mouse chromosomal map. Out of 24 new informative human colon cancer susceptibility loci 23 mapped to the vicinity of mouse *Sluc* loci, so with the previous data this revealed in total 27 clusters of colon-lung cancer susceptibility genes containing loci from human, mouse and rat^11^. Nine of these 27 clusters were < 2.0 cM long.

Additional GWAS^12,13^ detected 33 new human colon cancer susceptibility loci. Combining GWAS SNPs with Oncotarget markers^14^ revealed 18 new human lung cancer susceptibility loci detected in populations of European origin, and 5 lung cancer susceptibility genes were detected in populations of Asian descent^10^, all at significance <5 × 10^−8^. Remarkably, <u>all</u> these 23 new human lung cancer susceptibility loci^10,14^ were clustered with colon cancer susceptibility loci (Table 1). Moreover, the lung cancer susceptibility locus *VTI1A*^10^ was independently identified as a colon cancer susceptibility locus^15^.

**Table 1.**
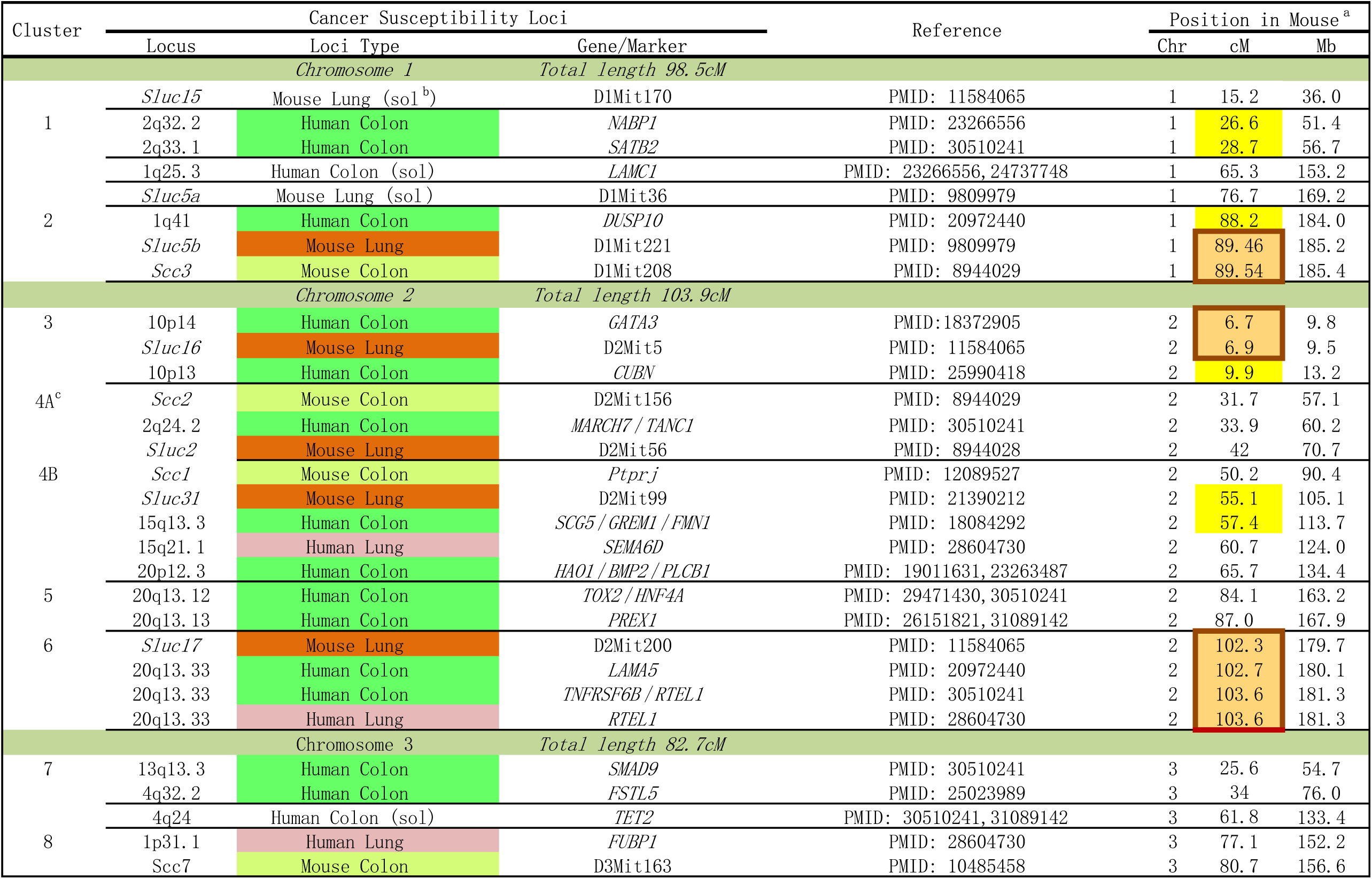
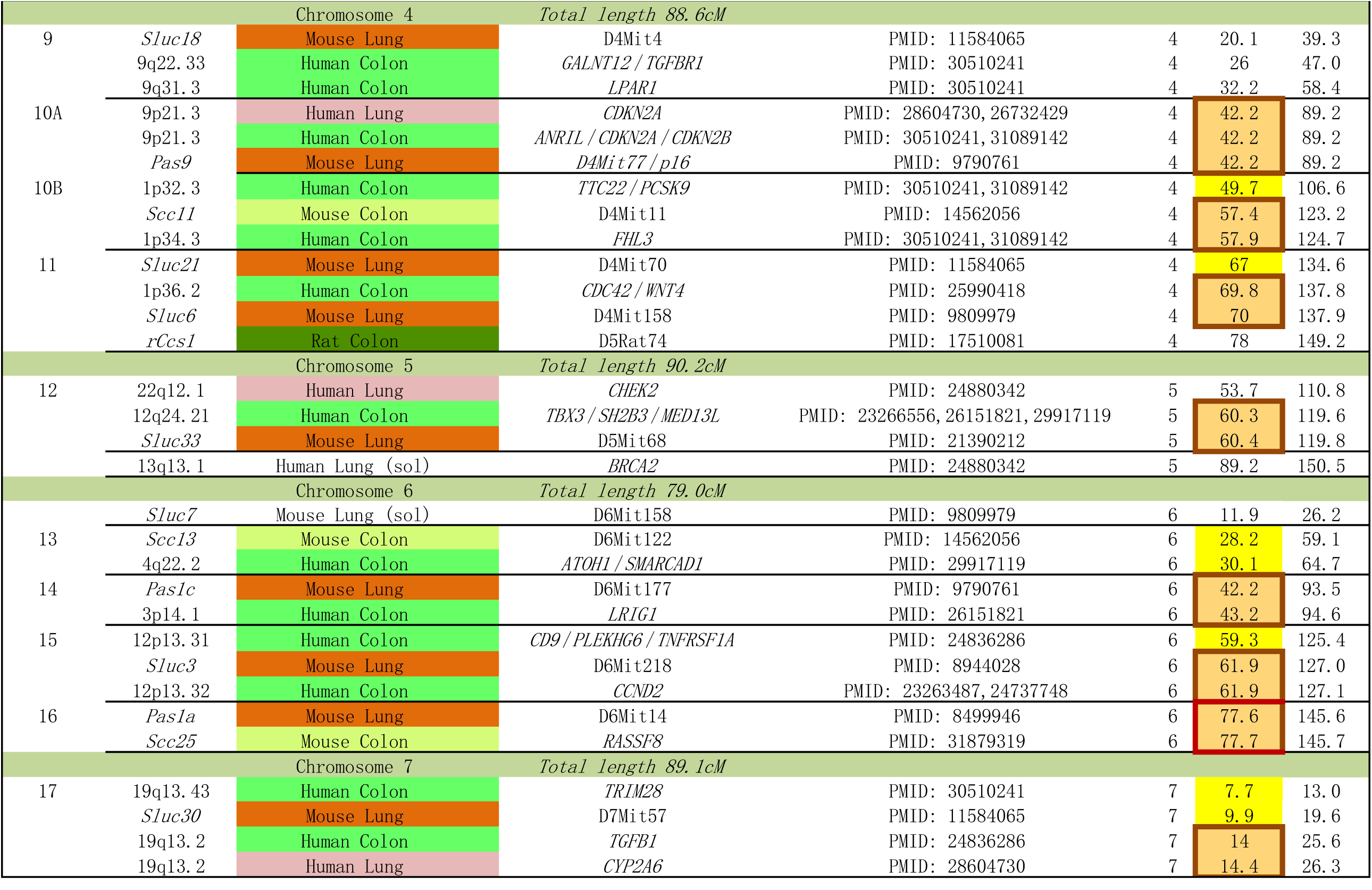
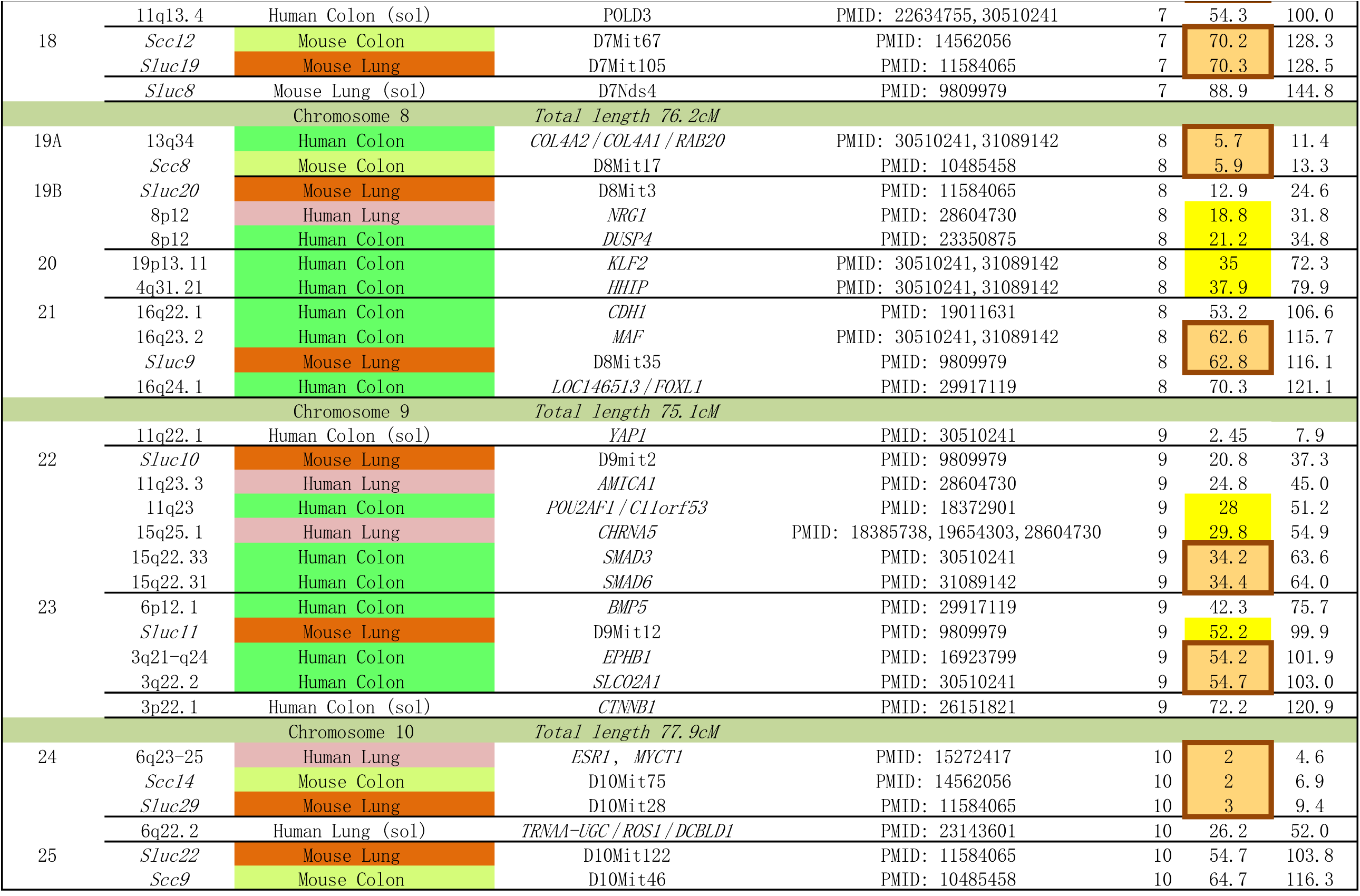
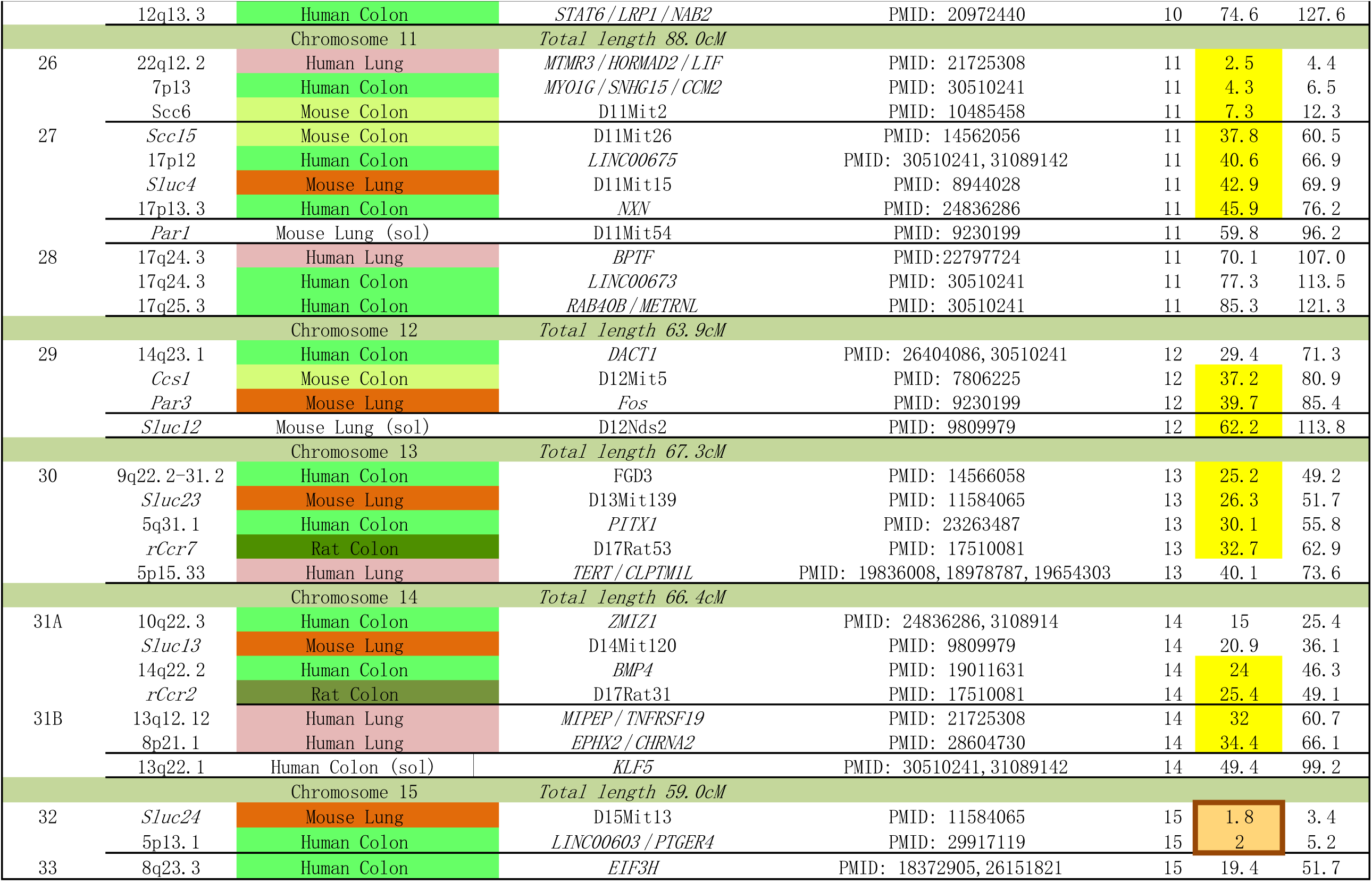
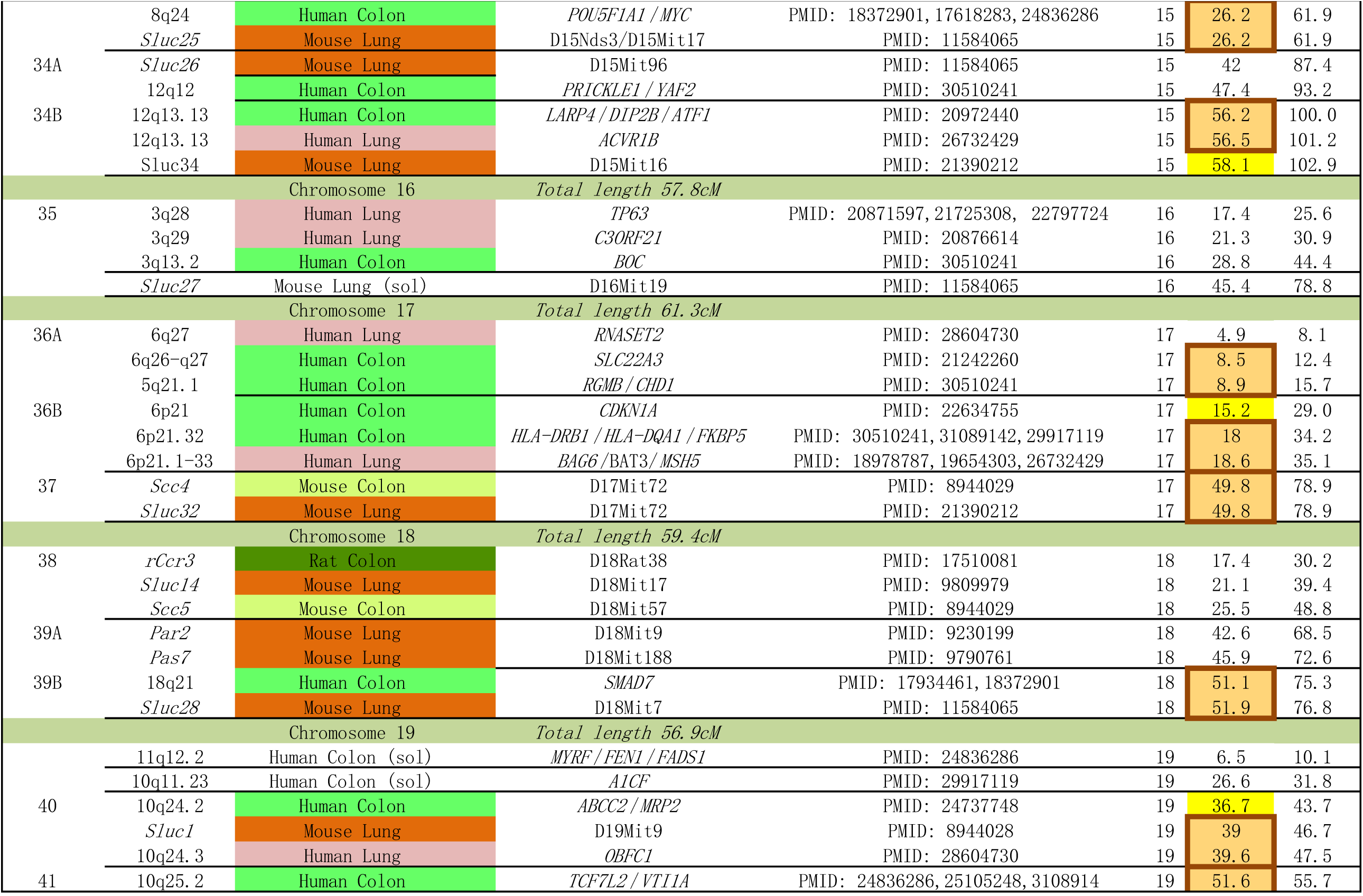
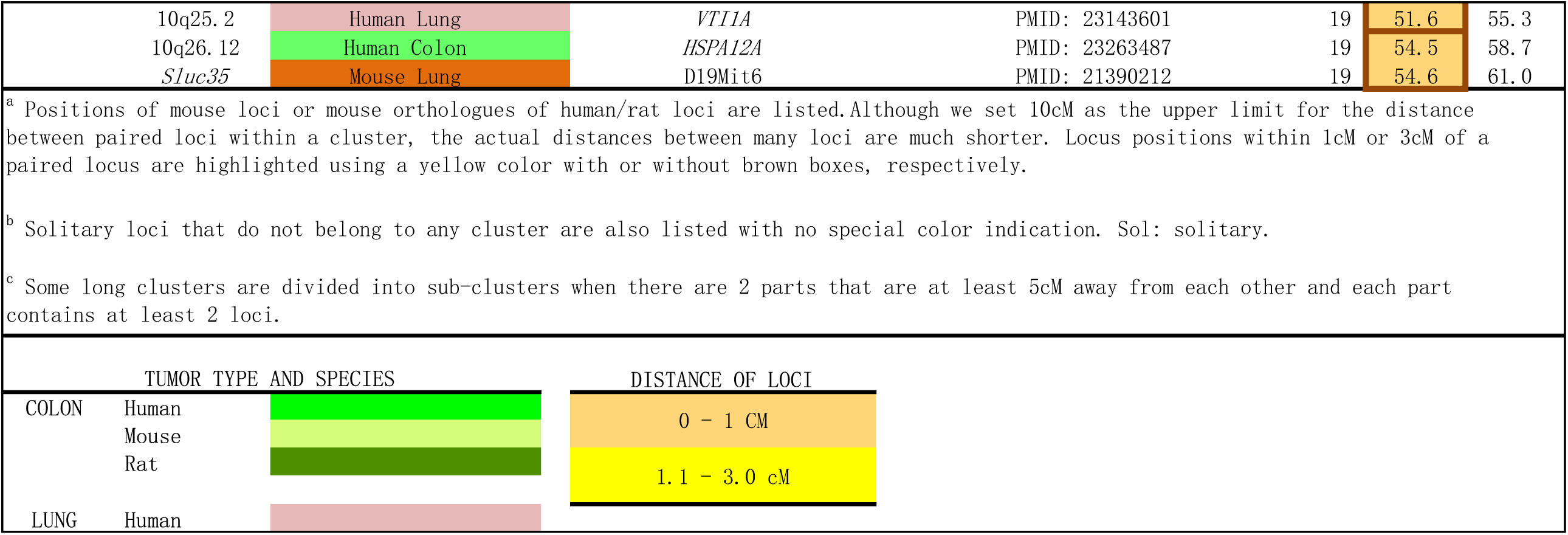
Co-localization of colon and lung cancer susceptibility loci identified in human, mouse and rat.

Collectively, this data comprised 159 chromosomal locations of colon and lung cancer susceptibility loci detected by linkage studies in humans, mouse, and rat and by GWAS in human. When their chromosomal locations were projected on the mouse genome, they formed 42 clusters (Table 1).

### Inter-species clustering

32 of the 42 clusters (76%) contained loci from at least two species (Table 1). This interspecies clustering likely reflects an extensive conservation of chromosomal positions of the colon and lung cancer susceptibility loci over >70 million years of separate evolution of mice, rats, and humans.

### Inter-organ clustering

37 of the 42 clusters (83.3%) contained both colon and lung cancer susceptibility loci (Table 1). Among the total of 159 analyzed loci, 77/95 colon cancer loci co-localized with at least one lung cancer locus: 22 of them within <1 cM and 18 within 1-3 cM; 56 of 64 lung cancer loci co-localized with at least one colon cancer locus: 22 of them within <1 cM and 12 within 1-3 cM. In order to investigate whether there is a preponderance of lung cancer loci in some clusters and colon cancer loci in others, we tested correlation of numbers of the two types of loci in the clusters that were defined in Table 1. If the two types of loci would be preferentially present in different clusters, this correlation would be negative. However, this was not the case, as this correlation was weakly but significantly positive (r=0.329, p=0.033).

### Statistical evaluation of colon-lung cancer susceptibility clustering

The most numerous clusters were the very tight ones: 54/159 loci were in clusters less than 1 cM long, which included 10 clusters shorter than 0.1 cM containing 21 loci, 8 clusters 0.1 – 0.5 cM long containing 18 loci,, and 7 clusters 0.5 - 1.0 cM long containing 15 loci. In contrast, the average distance of the 159 cancer susceptibility loci in the total autosomal mouse genome (1441 cM) was 9.06 cM. Only 16 of the 159 loci did not belong to any cluster and they were >10 cM distant from another colon or lung susceptibility locus.

The statistical analysis by Monte Carlo simulations examining the clustering of colon and lung cancer susceptibility loci using 100,000 simulated databases at window widths of 1, 2, 5, 10, 15, and 20 cM indicated that the p values of the deviation of the actual distribution of the colon and lung susceptibility loci from random distribution at these window widths were 0.00067, 0.093, 0.070, 0.056, 0.799, and 0.44, respectively. This agreed with the numerous short inter-specific clusters of colon and lung cancer susceptibility loci mentioned above and shown in Table 1.

## 3. Concordant RC strain susceptibility to colon and lung tumors

Due to different anatomical structure of colon and lung and different growth patterns of colon and lung tumors, the tumor parameters most affected by genetics are colon tumor numbers per mouse^3,5,8,11^ and lung tumor load per lung per mouse and lung tumor number per mouse^3,5,9^. The reason for it is that while mouse colon tumors mostly keep growing as separate structures, growing lung tumors fuse with neighboring tumors and form large confluent masses. Previously we have shown in two independent lung tumor induction experiments using Wilcoxon rank sums two sample tests that the the colon tumor susceptible strain CcS-19 has significantly higher lung tumor load than the colon tumor resistant strain CcS-20 (p<0.0001) and the colon tumor susceptible strain CcS-11 has a higher lung tumor load than the colon tumor resistant strain strain CcS-10 (p=0.0012) – see Fig. 2B in ref^5^, and the same difference caused by these strains in a separate experiment measuring lung tumor numbers (Support. Fig. 1 and Support. Table 1) in ref.^5^

To analyze comprehensively the RC strain differences in lung tumor susceptibility in relation to those in colon tumor susceptibility we compared the differences in lung tumor numbers caused by genotypes of the strains CcS-10, CcS-11, CcS-19 and CcS-20 by the t-distributed linear contrast test, that allows to test whether the strains CcS-10 and CcS-20 differ in lung tumor numbers from the strains CcS-11 and CcS-19.

### Results from the linear contrast analysis of the lung tumor count data

Data consist of tumor counts as outcome variable, with listing of strain (CcS-10, CcS-11, CcS-19, and CcS-20) and induced tumor type (lung).

The analysis of interest was to examine the specific contrast of testing for a difference between Strains (CcS-10, and CcS-020) versus Strains (CcS-11 and CcS-19), among the lung cancer induced mice. The results from testing this contrast are given here:

~~~
> summary(G, test=adjusted(‘single-step’))
Simultaneous Tests for General Linear Hypotheses
Multiple Comparisons of Means: User-defined Contrasts
Fit: lm(formula = TumorCts ∼ Strain, data = lungdat)
Linear Hypotheses:
c10c20vsc11c19 == 0, Estimate = -6.470, Standard error
1.718, t value= -3.765, Pr(>|t|) 0.000491 *** (44 degrees of
freedom)
---
 (Adjusted p values reported -- single-step method)
~~~

These results indicate that there is a significant value for the contrast, which would be interpreted as demonstrating that, among the mice treated to induce lung tumors, the two strains CcS-10 and CcS-20 differ significantly in tumor counts compared to the two strains CcS-11 and CcS-19.

These aren’t pairwise comparisons, but rather using the contrast formulation to aggregate the strains of interest to examine for significant differences in tumor counts.

### The independent genetic mapping of lung tumor susceptibility in a cross between strains highly susceptible and highly resistant to <u>colon</u> tumors revealed *Sluc/Pas* loci that co-localized with *Scc* loci

We tested whether the concordant resistance of CcS-10 to colon and lung tumors and concordant susceptibility of CcS-19 to these two tumors are caused by similar sets of loci. CcS-10 and CcS-19 are RC strains each containing a different set of 12.5% genes of the strain STS (highly susceptible to colon tumors) and 87.5% of BALB/c genes^3,5,16^. To determine whether the lung tumor susceptibility’s concordance with colon tumor susceptibility in these two strains is caused by related loci, we performed an independent linkage study of lung tumor susceptibility in F2 hybrids between the lung and colon tumor resistant strain CcS-10 and the lung and colon tumor susceptible strain CcS-19. We induced lung tumors in 226 (CcS-10 x CcS-19)F2 hybrids and by their genotyping we detected and mapped 7 *Sluc* loci with <u>significant main</u> effect on lung tumor size or load in these mice. Every detected *Sluc* locus was closely linked with a colon cancer susceptibility locus, either of mouse (Chr. 1 *Sluc-5b – Scc3*, Chr. 2 *Sluc31 – Scc1*, Chr.10 *Sluc29 - Scc14*, Chr. 18 *Sluc14 – Scc5*), or of human (Chr. 4 *Pas9* – 9p21.3, Chr. 5 *Sluc33* – 12q24.21, Chr. 8 *Sluc9 –* 16q24.1) (Fig. 1 and Table 1). No other significant loci were detected in this cross. This strongly suggests that in CcS-10 and CcS-19 the susceptibility to the two tumors is influenced by the effects of the paired colon and lung cancer susceptibility loci^5,11^.

## 4. Linkage of *DUSP* family of protein phosphatase genes with colon and lung cancer susceptibility loci

A surprising feature of the clusters of colon and lung cancer susceptibility genes reported here is their frequent and tight linkage with *DUSP* (*Dual specificity phosphatase*) genes (Table 2). Many DUSPs are closely linked to these clusters or located within them. DUSP enzymes remove phosphate groups from phosphorylated serine, threonine, and tyrosine from various proteins, thereby significantly modifying their functions^17^. *DUSP* genes encode six types of dual specificity phosphatases^17^ and the close linkage with colon and lung cancer susceptibility genes applies to all of them. Table 2 shows their chromosomal location in the mouse and their distance in cM from the projected location of the nearest mouse, rat, or human colon or lung cancer susceptibility locus (as listed in Table 1). DUSP enzymes have manifold effects in development and progression of cancer^18,19,20^ that include also modulation of effects of mitogen induced kinases.Out of 45 *DUSP* genes, 13 (29%) are within less than 1 cM of a colon or lung cancer susceptibility locus, 20 (44%) are within less than 2 cM of such locus, and 33 (73%) within less than 5 cM of such locus. Several of them are located inside the clusters of colon-lung cancer susceptibility loci. This very close proximity of many *DUSP* genes with colon or lung cancer susceptibility genes suggests that some of them may be candidates for these susceptibility genes. In fact, some *DUSP* genes have been directly implicated as cancer susceptibility genes. *DUSP10* has been identified in GWAS as a human colon cancer susceptibility gene and it is 1.2 – 1.4 cM distant from the mouse colon and lung cancer susceptibility loci *Scc3* and *Sluc5b*^21^, respectively. *DUSP4* has also been suggested in an early GWAS^22^ as a potential candidate gene for human colon cancer susceptibility gene at 8p12, and although falling short of the standard significance threshold, it co-localizes also with a human lung cancer susceptibility gene at 8p12 in cluster 19 (Table 1). The *DUSP* family member *PTEN* has tumor suppressor effects and it is the second most frequently deleted gene in cancers^17^.

**Table 2.**
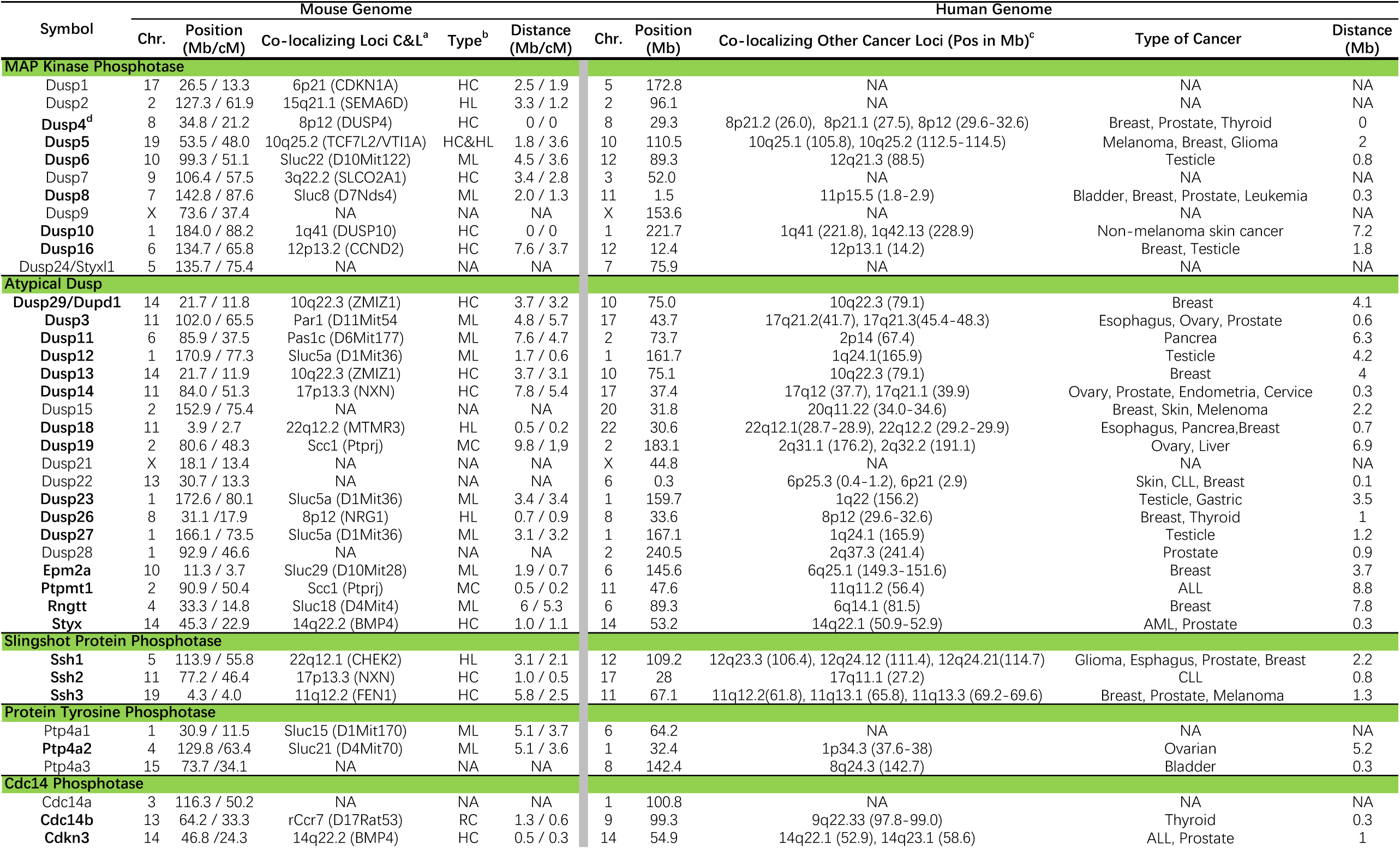
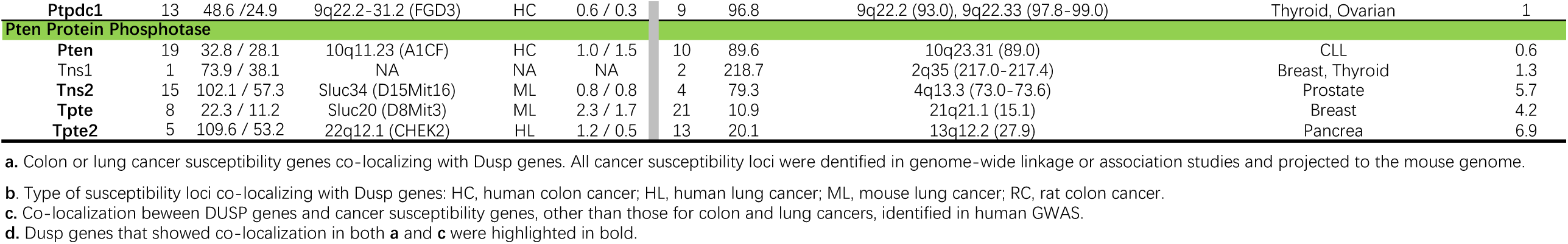
Dusp gene family and cancer susceptibility genes.

We evaluated statistically the tendency of *DUSP* genes to occur in proximity to the colon/lung susceptibility loci via Monte Carlo Simulation methods. Specifically, we used simulation to investigate whether the actual position of *DUSP* genes vis-à-vis the susceptibility loci were in closer proximity than would be expected if the same *DUSP* loci occurred in random positions in the genome. We used the arithmetic mean distances of simulated random *DUSP* positions to the nearest colon-lung susceptibility locus and compared them to the true mean distances of *DUSP* genes to the nearest colon-lung susceptibility locus. We found that the true *DUSP* positions are on average closer to colon-lung susceptibility loci than would be expected by random chance, with empirical p-value of 0.02, based on 100,000 simulated datasets. In summary, these analyses show that *DUSP* genes are significantly more likely to occur in close proximity to colon or lung loci than would be expected by chance, and that the true mean values of distances of *DUSP* genes to susceptibility loci for colon and lung cancer are substantially below those means in a collection of randomly distributed *DUSP* positions.

As 7 out of 45 *DUSP* genes (2 of them located on X chromosome) were not linked to any of the colon or lung susceptibility loci, we investigated whether these and other D*USP* genes were linked to susceptibility genes for cancers of other organs using their map positions reported in ref^4^. As shown in the right-hand half of the Table 2, with the exception of the two X-linked loci *DUSP9* and *DUSP21* and *CDC14A* (on human chromosome 1), every *DUSP* gene is linked to at least one cancer susceptibility gene. A possible corollary of this observation is that many of the susceptibility genes for cancers in other organs map close to the colon-lung cancer susceptibility clusters.

## Discussion

We highlighted here four previously non-reported aspects of genetics of colon and lung cancer susceptibility: significant pairwise clustering of susceptibility loci for colon and lung cancer, its conservation in three species for >70 million years, correlated direction of phenotypic effect of the paired loci on both colon and lung tumorigenesis in recombinant congenic strains, and frequent association of *DUSP* genes with colon and lung cancer susceptibility genes. These data offer a possibility to extend the understanding of individual genetic susceptibility to colon and lung cancers, its molecular and functional basis, and the responsible genes, which at the present are not sufficiently defined.

The data from three species that strongly indicate an evolutionary conserved co-localization of most colon and lung cancer susceptibility genes and the concordant direction of effects of the paired loci is compatible either with a single susceptibility gene or shared conserved regulatory elements, although more complex explanations will likely emerge too.

The precision of map positions of genes in Table 1 may be negatively affected by mapping the same gene in different studies and different species using different but close DNA markers and by intra- and inter-species indel polymorphisms. Moreover, many SNPs most significantly associated with QTLs are close to genomic regulatory elements and it is conceivable that different elements may control the same gene in different organs (or species). Finally, some cancer QTLs represent complex of interacting genes and or regulatory elements spanning several megabases of genome (reference^4^, Figure 3). These complications, however, do not invalidate the evidence for clustering, as the length of inter-cluster intervals far exceeds the potential mapping imprecisions. To the contrary, this suggests that many members of a cluster may be more tightly linked than the Table 1 shows. Sud et al. ^4^4 reported that a sizeable proportion of SNPs is associated with multiple cancers and we showed here that the pair-wise association of colon and lung cancer susceptibility loci is evolutionary conserved and hence likely based on shared functional pathways, most of which are not yet known. We did not investigate systematically whether a similarly strong clustering of susceptibility loci for other tumor types, but data in Table 2 suggest that it may be detected later. A shared heritability of six solid cancers was indicated by meta-analysis of GWAS data^23^, but no information about the loci involved was presented. Our experimental design in mice^3,5^ permits in the future to assay separately the effects of different alleles of individual loci on different aspects of colon and lung cancer susceptibility in terms of quantitative and qualitative tumor pathology, their transcriptional and epigenetic effects in the target organs and tumors (for example^24,25^), somatic mutations^26^ and inflammatory response^27^. Results of such studies can be further expanded by targeted manipulation of candidate susceptibility genes. Analyzing effects of single susceptibility genes in mice could help to disentangle the control of individual colon and lung cancer susceptibility in humans as well. In addition, close association of some of them with *DUSP* genes can help to understand the mechanisms of their effects.

The inter-species conservation of these “early” cancer QTLs suggests that they likely operate in important biological pathways.Their high replicability makes them potentially useful for understanding the mechanisms of individual genetic risk of these two frequent cancers and for development of novel preventative and therapeutic methods. Recent high-powered GWAS meta-analyses in cancer and other diseases identified very large numbers of novel QTLs. It is possible that with numbers of QTLs for each tumor type exceeding several hundreds, their ensuing high density in genome can make it difficult to detect statistically their systemic colocalization. However, the future contribution of these new QTLs to personal medicine requires that they exhibit genetic stratification^28^, but until now the predominant contribution to genetic risk has been provided mainly by the early detected QTLs, because they tend to have larger phenotypic effects^29,30^. A study of effect-size distribution of susceptibility QTLs of 14 cancers suggests that in spite of considerable quantitative differences between cancer types, the cumulative effects of stronger and thus “earlier” QTLs will likely determine the largest part of individual genetic susceptibility^31^. The present study is based on such “medium strength” QTLs that were discovered in the early GWAS involving a relatively limited number of cases and controls^10,12,13,14^ compared to the recent meta-analyzes. All GWAS-detected human cancer susceptibility loci analyzed here were described in the early genome-wide association studies^10,12,13,14^ in which they were identified with the required significance level of 5 × 10^−8^. No loci from other GWAS were included. Such early detected loci appear to be exceptionally replicable, because they were not only detected in the early GWAS, but also were over-represented among the loci detected in all subsequent GWAS due to their repeated detection^29^. Moreover, generally the overall disease predictive capacity of GWAS loci, measured as area under the ROC curve (AUC), tended to increase with the number of loci detected in the early GWAS, but addition of larger numbers of loci first detected in later GWAS is predicted to result in a much smaller improvement of overall disease predictive capacity^29,30^. This implies that the human paired colon-lung susceptibility loci analyzed in the present study, which were all detected in the early GWAS studies, will also likely remain responsible for a relatively large proportion of genetic risk for these two cancers.

## Ethics Statement

All animal experiments were approved by the IACUC committee at Roswell Park Cancer Institute (IACUC protocol 905M).

## Mice

Mice received acidified drinking water (pH 2.5–3.0) and a standard laboratory diet (LM-485, Harlan Teklad, U.S.) ad libitum. Recombinant congenic (RC) strains are sets of 20 inbreed strains derived from the same parental strains. Each CcS RC strain (CcS-1 through CcS-20) has 87.5% of the genome from the “background” BALB/c strain and 12.5% of the genome from the “donor” STS strain^3,5^ Each OcB RC strain (OcB-1 through OcB-20) has 87.5% of the genome from the “background” O20 strain and 12.5% of the genome from the “donor” B10.O20 strain.

## Tumor Induction and Histological Analysis

For lung tumor induction, pregnant female mice were given an intraperitoneal (i.p.) injection of 30 mg/kg body weight of the carcinogen N-ethyl-N-nitrosourea (ENU) dissolved in phosphate-buffered citric acid (pH 5.8) at day 17 of gestation. The offspring of carcinogen-injected females were thus exposed to ENU transplacentally and their whole lungs were examined for tumors at the age of four months. Colon tumors were induced in adult mice with eight weekly subcutaneous injection of 15 mg/kg body weight of azoxymethane (AOM) or 1,2-dimethylhydrazine (DMH). Mice were euthanized after four months since the last injection and their colons were examined. All tissues were removed, fixed in 10% neutral buffered formalin and embedded in histowax. The embedded lungs and colons were sectioned semi-serially (5-mm sections at 100-mm intervals), stained with haematoxylin-eosin and examined microscopically at 50x and 400x magnifications. To distinguish unequivocally individual tumors, position of a tumor in sequential sections, its shape and size, positional relation to bronchi or blood vessels, and other characteristics of tumor cells have been used.

## Genotyping

More than 90 % of the genetic material from the ‘donor’ strain in a RC strain is concentrated in 9 to 13 discrete contiguous chromosomal regions with intermediate length (5–25 cM) that are usually located on 7 to 11 different chromosomes^3^. The positions and lengths of the majority of the ‘donor’ strain-derived chromosomal regions in CcS and OcB RC strains have been determined with more than 500,000 microsatellite or single nucleotide polymorphism (SNP) markers across the whole genome. Based on such information, each known segregating chromosomal region is represented by at least one genetic marker. More markers have been tested in the longer donor chromosomal regions and the maximal distance between two markers was less than 10 cM.

## Definition of Clusters

Susceptibility loci for colon and lung cancers were retrieved from literature and the NHGRI GWAS catalog (for human GWAS loci only). Only loci with a genome-wide significance were retained, i.e. p<5×10^−8^ for GWAS loci in human or P<0.05 after Bonferroni correction of multiple testing for loci in rat and mouse. Entries for which a genomic region and/or association statistic could not be unambiguously assigned were excluded. Loci identified in human and rat were projected to the mouse genome based on positions of their orthologous regions in mice, acquired from the major genome browsers, i.e. the Jackson Laboratory (https://informatics.jax.org), the Rat Genome Database (https://rgd.mcw.edu), Ensemble (https://www.ensembl.org) and the UCSC genome browser (https://www.genome.ucsc.edu). Two or more loci are defined as a cluster when their distance is less than 10 cM. Although we set 10cM as the upper limit for the distance between paired loci within a cluster, the actual distances between most loci in a cluster are much shorter (Table 1).

## Statistical analysis

### Concordance of strain differences in colon and lung tumor susceptibility

Due to different anatomical structure of colon and lung and different growth patterns of colon and lung tumors, the tumor parameters most affected by genetics are colon tumor numbers per mouse^3,5,8,11^ and lung tumor load per lung per mouse^3,5,9^. The reason for it is that while mouse colon tumors mostly keep growing as separate structures, growing lung tumors fuse with neighboring tumors and form large masses. To compare the quantitative variation of the two phenotypes in the strains CcS-10, CcS-11, CcS-19 and CcS-20, we ranked all AOM-treated individual mice for number of colon tumors and all individual ENU treated progeny for lung tumor load. These rank-normalized values in different strains were compared by the t-distributed linear contrast test at 44 degrees of freedom.

### Assessment of clustering

Monte Carlo multi-window analysis was used to assess the distribution of positions of the mouse colon and lung cancer susceptibility loci in mouse autosomal genome and the orthologous positions of human and rat colon and lung cancer susceptibility loci projected on the mouse autosomal genome as listed in Table 1. We accounted for chromosome length, and only included within-chromosome distances. The test involved five window widths (1, 2, 5, 10, and 20 cM) and 100.000 simulations for each of them.

Similarly, Monte Carlo analysis of closeness of *DUSP* loci to colon/lung susceptibility loci. We took the true positions of the susceptibility loci, and the true positions of the *DUSP* loci, and computed the set of pairwise distances of a *DUSP* locus with its nearest susceptibility locus, and computed the arithmetic means of this collection of true proximity values. We then generated a random set of the same number of *DUSP* positions across the genome and computed the set of pairwise distances of a random *DUSP* position with its nearest susceptibility locus, and computed means of this set of distances. We accounted for chromosome length, and only included within-chromosome distances in determining nearest locus to a *DUSP* position. We repeated the random set generation 100,000 times and computed the number of these random datasets that had mean value below the true means, using both arithmetic and geometric means, which provided empirical p-values for comparing true mean *DUSP* proximities to an equivalent mean of random positions.

## Acknowledgements

We thank Dr. Alan Hutson, Chairperson of Department of Biostatistics and Bioinformatics of the Roswell Park Comprehensive Cancer Center for generous support, discussions and advice.

This work had been supported by the NCI grant R01 CA 115158 and an institutional grant from the Roswell Park Comprehensive Cancer Center to P. Demant, and by NCI grant P30CA016056 to the Roswell Park Comprehensive Cancer Center supporting the use of its Biostatistics and Genomic Shared Resources and by the grant 16JCYBJC from the Natural Science Foundation from the Tianjin City and Tianjin Thousand Young Talents Plan to L. Quan.

## References

1. Pharoah, P. D. et al. Polygenic susceptibility to breast cancer and implications for prevention. Nat Genet 31, 33–36, doi:10.1038/ng853 (2002).

2. Balmain, A., Gray, J. & Ponder, B. The genetics and genomics of cancer. Nat Genet 33 Suppl, 238–244, doi:10.1038/ng1107 (2003).

3. Demant, P. Cancer susceptibility in the mouse: genetics, biology and implications 021521for human cancer. Nat Rev Genet 4, 721–734, doi:10.1038/nrg1157 (2003).

4. Sud, A., Kinnersley, B. & Houlston, R. S. Genome-wide association studies of cancer: current insights and future perspectives. Nat Rev Cancer 17, 692–704, doi:10.1038/nrc.2017.82 (2017).

5. Quan, L. et al. Most lung and colon cancer susceptibility genes are pair-wise linked in mice, humans and rats. PLoS One 6, e14727, doi:10.1371/journal.pone.0014727 (2011).

6. De Miglio, M. R. et al. Identification and chromosome mapping of loci predisposing to colorectal cancer that control Wnt/beta-catenin pathway and progression of early lesions in the rat. Carcinogenesis 28, 2367–2374, doi:10.1093/carcin/bgm119 (2007).

7. Bissahoyo, A. C. et al. A New Polygenic Model for Nonfamilial Colorectal Cancer Inheritance Based on the Genetic Architecture of the Azoxymethane-Induced Mouse Model. Genetics 214, 691–702, doi:10.1534/genetics.119.302833 (2020).

8. Chen, J. & Huang, X. F. The signal pathways in azoxymethane-induced colon cancer and preventive implications. Cancer Biol Ther 8, 1313–1317, doi:10.4161/cbt.8.14.8983 (2009).

9. Festing, M. F. et al. At least four loci and gender are associated with susceptibility to the chemical induction of lung adenomas in A/J x BALB/c mice. Genomics 53, 129–136, doi:10.1006/geno.1998.5450 (1998).

10. MacArthur, J. et al. The new NHGRI-EBI Catalog of published genome-wide association studies (GWAS Catalog). Nucleic Acids Res 45, D896–D901, doi:10.1093/nar/gkw1133 (2017).

11. Quan, L., Hutson, A. & Demant, P. Cross-Cancer Analysis Reveals Novel Pleiotropic Associations-Letter. Cancer Res 77, 6042–6044, doi:10.1158/0008-5472.CAN-16-3262 (2017).

12. Huyghe, J. R. et al. Discovery of common and rare genetic risk variants for colorectal cancer. Nat Genet 51, 76–87, doi:10.1038/s41588-018-0286-6 (2019).

13. Law, P. J. et al. Association analyses identify 31 new risk loci for colorectal cancer susceptibility. Nat Commun 10, 2154, doi:10.1038/s41467-019-09775-w (2019).

14. McKay, J. D. et al. Large-scale association analysis identifies new lung cancer susceptibility loci and heterogeneity in genetic susceptibility across histological subtypes. Nat Genet 49, 1126–1132, doi:10.1038/ng.3892 (2017).

15. Wang, H. et al. Trans-ethnic genome-wide association study of colorectal cancer identifies a new susceptibility locus in VTI1A. Nat Commun 5, 4613, doi:10.1038/ncomms5613 (2014).

16. Moen CJ, et al. The recombinant congenic strains--a novel genetic tool applied to the study of colon tumor development in the mouse. Mamm Genome 1, 217–27. (1991)

17. Patterson, K. I. et al. Dual-specificity phosphatases: critical regulators with diverse cellular targets. Biochem. J. 418, 475–489, doi: 10.1042/bj20082234 (2009).

18. Keyse, S. M. Dual-specificity MAP kinase phosphatases (MKPs) and cancer. Cancer Metastasis Rev 27, 253–261, doi:10.1007/s10555-008-9123-1 (2008).

19. Meeusen, B. & Janssens, V. Tumor suppressive protein phosphatases in human cancer: Emerging targets for therapeutic intervention and tumor stratification. Int J Biochem Cell Biol 96, 98–134, doi:10.1016/j.biocel.2017.10.002 (2018).

20. Ruvolo, P. P. Role of protein phosphatases in the cancer microenvironment. Biochim Biophys Acta Mol Cell Res 1866, 144–152, doi:10.1016/j.bbamcr.2018.07.006 (2019).

21. Png, C. W. et al. DUSP10 regulates intestinal epithelial cell growth and colorectal tumorigenesis. Oncogene 35, 206–217, doi:10.1038/onc.2015.74 (2016).

22. Fernandez-Rozadilla, C. et al. A colorectal cancer genome-wide association study in a Spanish cohort identifies two variants associated with colorectal cancer risk at 1p33 and 8p12. BMC Genomics 14, 55, doi:10.1186/1471-2164-14-55 (2013).

23. Jiang, X. et al. Shared heritability and functional enrichment across six solid cancers. Nat Commun 10, 431, doi:10.1038/s41467-018-08054-4 (2019).

24. Dampier, C.H. et al. Oncogenic Features in Histologically Normal Mucosa: Novel Insights Into Field Effect From a Mega-Analysis of Colorectal Transcriptomes. Clin Transl Gastroenterol 11, e00210 (2020).

25. Diez-Obrero, V. et al. Genetic Effects on Transcriptome Profiles in Colon Epithelium Provide Functional Insights for Genetic Risk Loci. Cell Mol Gastroenterol Hepatol 12, 181–197 (2021).

26. Rogers, Z.N. et al. Mapping the in vivo fitness landscape of lung adenocarcinoma tumor suppression in mice. Nat Genet 50, 483–486 (2018).

27. Zhang, Y. D. et al. Assessment of polygenic architecture and risk prediction based on common variants across fourteen cancers. Nat Commun 11, 3353, doi:10.1038/s41467-020-16483-3 (2020).

28. R. Lang, M. Hammer, J. Mages, DUSP meet immunology: dual specificity MAPK phosphatases in control of the inflammatory response. J Immunol 177, 7497–7504 (2006).

29. Wray, N. R., Wijmenga, C., Sullivan, P. F., Yang, J. & Visscher, P. M. Common Disease Is More Complex Than Implied by the Core Gene Omnigenic Model. Cell 173, 1573–1580, doi:10.1016/j.cell.2018.05.051 (2018).

30. Marigorta, U. M., Rodriguez, J. A., Gibson, G. & Navarro, A. Replicability and Prediction: Lessons and Challenges from GWAS. Trends Genet 34, 504–517, doi:10.1016/j.tig.2018.03.005 (2018).

31. Turnbull C., Sud A., Houlston R. C. Cancer genetics, precision prevention and a call for action. Nature Genet. 50: 1212–1218: doi:10.1038/s41588-018-0202-0 (2018).

32. Zhang Y. D. et al. Assessment of polygenic architecture and risk prediction based on common variants across common cancers. Nat Commun 11, 3353, doi: 10.1038/s41467-020-16483-3 (2020).

